# COMT inhibitor tolcapone represses the migration and invasion of trophoblast and results in preeclampsia-like phenotypes in mice with the modulation of EMT signaling

**DOI:** 10.1101/2023.03.17.530517

**Authors:** Lu Chen, Yuexia Li, Menglan Pang, Qiuling Jie, Fei Sun, Liping Huang, Yanlin Ma, Fei Wu, Xiaojing Yue

## Abstract

Pregnancy in young-onset Parkinson’s disease (YOPD) is rare; however, medication for this condition is critical for maternal and fetal health. Tolcapone is an effective antiparkinsonian drug. However, its safety studies on mothers and fetuses are limited. In this study, we aimed to investigate the effects of tolcapone on the mother and the developing fetus during different stages of gestation. Tolcapone was administered to pregnant mice at a dose of 60 (H-Tol) or 30 mg/kg/day (L-Tol) during early or mid-gestation. We observed that tolcapone administration during early gestation causes abortion and delays fetal development in a dose-dependent manner. During mid-gestation tolcapone barely caused embryo lethality; however, the mice developed preeclampsia-like phenotypes, including maternal hypertension, proteinuria, and fetal growth restriction. Histomorphological analysis of placentas from tolcapone treated mice revealed abnormalities in the trophoblast layer and the impaired trophoblast invasion in the decidua. Mechanistically, we revealed that tolcapone inhibits the invasion and migration of trophoblasts *in vitro*, with changes in the protein expression of Snail, Twist, and E-cadherin. In conclusion, tolcapone caused embryo lethality and growth restriction during early gestation, whereas it caused preeclampsia-like phenotypes in mice with defective trophoblast invasion in mid-gestation. Collectively, our study provides novel insights into the effects of tolcapone on pregnancy.

## Introduction

Young onset Parkinson’s disease (YOPD) is a type of Parkinson’s disease (PD) diagnosed between 21 and 40 years[1]. The YOPD population comprises 5-7% of patients with PD in Western countries and approximately 10% in Japan[2]. Unlike late-onset PD, YOPD progresses slowly with regular and continuous care and treatments[3]. Pregnancy in YOPD is rare; however, clinicians should counsel women with YOPD of childbearing potential. Based on an observation of 74 pregnant women with PD, antiparkinsonian medications improved PD symptoms and reduced the incidence of deteriorated symptoms from 67% to 36%[4]. Antiparkinsonian medications are effective for controlling PD in pregnancy; however, knowledge of the safety of medications during pregnancy and the potential risk to the fetus remains limited[5].

Tolcapone, an inhibitor of catechol O-methyltransferase (COMT), is indicated in combination with levodopa/carbidopa therapy for PD[6–8]. The Food and Drug Administration (FDA) lists antiparkinsonian drugs in pregnancy category C[9]. Pregnancy category C implies that animal studies revealed a risk to the fetus, and studies on humans are unavailable. The potential benefits in maternal disease control may outweigh the potential risks to the fetus. Administration of tolcapone during animal gestation has been implicated in fetal growth restriction and malformation; however, pathological change in placenta remains unknown and potential adverse effect on mother is not adequately studied[10].

The placenta is vital for evaluating the tolcapone-mediated risks to the fetus. Notably, changes in placenta, which is an important tool for understanding the mechanisms of reproductive and developmental toxicity, must be evaluated[11]. As the main cell type in the placenta, trophoblasts anchor the fetus to the uterine wall, synthesize steroids and hormones, and exchange nutrients and gases between mother and fetus. Trophoblast cell invasion into the maternal decidua is critical for remodeling the uterine spiral artery into a dilated vessel. Failure of invasive trophoblasts can lead to failed spiral artery transformation and placental ischemia, with consequences for fetal growth restriction and other pregnancy complications such as preeclampsia[12–14]. Epithelial-mesenchymal transition (EMT) signaling is pivotal in mediating trophoblast invasion. Several *in vitro* experiments indicated that cell adhesion molecule E-cadherin hampers trophoblast invasion and migration, while the upstream transcription factors Snail and Twist promote trophoblast invasion[15]. (Twist modulates human trophoblastic cell invasion via regulation of N-cadherin; Activin A Increases Human Trophoblast Invasion by Inducing SNAIL-Mediated MMP2 Up-Regulation Through ALK4) Upregulation of E-cadherin and downregulation of Snail were noted in human placentae with preeclampsia[16, 17]. (Down-regulation of the transcription factor snail in the placentas of patients with preeclampsia and in a rat model of preeclampsia) In the present study, we found that tolcapone inhibited trophoblast invasion with change in EMT signaling; mice developed preeclampsia-like phenotypes with mid-gestation tolcapone administration. Our study provides additional information of tolcapone in maternal-fetal and placental toxicity.

## Methods

### Animals and experimental protocol

C57BL/6 mice aged 8-10 weeks were housed overnight, and the presence of vaginal spermatozoa confirmed pregnancy. Pregnant mice on gestational day 0.5 or day 9.5 were administered tolcapone (A4383, APExbio, USA) once daily via gastric gavage until the end of gestation. A high dose (H-Tol, 60 mg/kg/day) of tolcapone for mice was derived from the daily recommended human dose (100-200 mg, thrice a day)[8, 18]. The low dose (L-Tol) of tolcapone administered to the mice was 30 mg/kg/day.

Blood pressure was measured with a Mouse Blood Pressure System (BP2010, Softron Biotechnology, China) using the tail-cuff method. The mice were trained for 2-3 days until they were accustomed to the cone animal holder and remained calm during the measurements. Blood pressure was recorded from gestational day 0-17.5.

All animal experiments were approved by the Institutional Animal Ethical Care Committee of Southern Medical University Experimental Animal Center.

### Cell culture

The human first-trimester placental trophoblast cell line HTR8/SVneo (ATCC CRL-3271, USA) was cultured in RPMI 1640 medium containing 2 mM glutamine, 10% FBS, and 1% penicillin-streptomycin. The human choriocarcinoma trophoblast cell line BeWo (ATCC CCL-98, USA) was cultured in F-12K medium containing 2 mM glutamine, 10% FBS, and 1% penicillin-streptomycin. The cells were maintained at 37°C with 5% CO_2_.

### Wound healing, Transwell, cell viability, and cell proliferation assays

Wound healing and Transwell assays were performed using the invasive trophoblast cell line HTR8/SVneo. The wound healing assay was conducted using an ibidi culture silicone insert (81176, ibidi, Germany). HTR8/SVneo cells were cultured to 90% confluence, and the insert was removed. The cells were maintained in a medium containing tolcapone and 1% FBS to minimize cell proliferation. Cell migration was examined after 24 h and 48 h. For Transwell migration assay, tolcapone-treated HTR8/SVneo cells were suspended in a serum-free medium and seeded into a Transwell chamber with 8μm pores (353097-8EA, Corning, USA). Medium containing 10% FBS was supplied to the lower chamber to induce cells to pass through the Transwell pores. The Transwell invasion assay was performed using a Transwell chamber pre-coated with Matrigel (354234, Corning, USA). After 24 h of incubation at 37℃ with 5% CO_2_, the cells on the lower surface of the transwell membrane were fixed and labeled with crystal violet dye. Images were acquired using a BX51 microscope (Olympus, Japan). Twenty stained images from each group were quantified using Image-J software (NIH, USA).

A cell viability assay was performed using HTR8/SVneo cells. The cells were exposed to 0-350 mM tolcapone for 24 h. Viable cells were assessed using the CCK-8 assay (C0039, Beyotime, China). For the cell proliferation assay, HTR8/SVneo cells were cultured in a medium containing 0-45 mM tolcapone for 72 h. The numbers of live cells were determined using the CCK-8 assays, and proliferation rates were analyzed.

### Western immunoblotting

Protein samples for western blotting were extracted as described previously[19]. Briefly, proteins were extracted from homogenized mouse placental tissues or cell samples using RIPA lysis buffer (89900, Thermo Fisher Scientific, USA) containing protease and phosphatase inhibitors (A32961, Thermo Fisher Scientific, USA). Equal protein loading was confirmed using the intensity of the GAPDH blot. Western blotting experiments were performed in triplicate. Densitometry of the blots was performed using Image-J software (NIH, USA).

The primary antibodies used in this study were as follows: anti-E-cadherin (3195), anti-Twist1 (31174), and anti-Snail (3879) from Cell Signaling Technology, USA, and anti-GAPDH (sc-47724) from Santa Cruz, USA.

### H&E staining, immunohistochemistry staining, and fluorescence staining

Paraffin-embedded sections of the whole mouse placentas were stained with hematoxylin and eosin staining (60524ES60, Yeasen, China). Consecutive mouse placental sections were used for immunohistochemical staining of cytokeratin 8 (CK8) and α-smooth muscle actin (α-SMA). The primary antibodies anti-CK8 (ab154301) and anti-α-SMA (ab5694) were from Abcam, USA. BrdU staining (6813S, Cell Signaling Technology, USA) of proliferating HTR8/SVneo and BeWo cells was performed. Nuclei were visualized by DAPI staining. Images were acquired using a fluorescence microscope (BX51 System Microscope, Olympus, Japan). Twenty stained images from each group were quantified using Image-J Software (NIH, USA).

### Statistical analysis

The data are expressed as the mean ± standard deviation (SD). Unpaired, two-tailed Student’s *t*-test was used for two-group comparisons. One-way ANOVA, followed by the Student–Newman–Keuls method, was used for multiple-group comparisons. All analyses were conducted using GraphPad Prism 8.0 and SAS 9.4. Statistical significance was set at *P*<0.05.

## Results

### Tolcapone causes abortion, fetal growth restriction and placental abnormality in early gestation

Administration of tolcapone in pregnant mice started at gestational day 0.5 and lasted until gestational day 17.5 (Fig. 1A). The dose of 60 mg/kg/day of tolcapone in mice was derived from the recommended daily human dose of 100-200 mg, thrice a day, which was set as the highest dose (H-Tol) in our animal study. To explore the minimum safe dose of tolcapone during pregnancy, we administered a low dose (L-Tol) of 30 mg/kg/day tolcapone. The body weights of L-Tol mice increased as gestation progressed; however, they were significantly lower from gestational day 7.5 to the end of gestation compared with those of the saline control group (Fig. 1B). The H-Tol mice displayed no change in body weight, indicating abortion or embryo loss (Fig. 1B). Therefore, we analyzed the placentas and fetuses of L-Tol mice in subsequent studies. The fetal size decreased at gestational day 17.5 in the L-Tol group compared with that in the saline control group (Fig. 1C). The number of live pups decreased, and the fetal resorption rate increased in L-Tol mice (Fig. 1D). Consistent with the smaller fetal size, the fetal weight of L-Tol mice decreased; however, placental weights increased (Fig. 1E). To explore the underlying mechanism, we examined the placental structure. Histomorphological analysis revealed a pathological change in the placentas of L-Tol mice, as collagen deposits were observed in the junction zone (JZ). Endothelial hyperplasia with thrombus formation was observed in large blood vessels (Fig. 1F). The ratio between the labyrinth zone (LZ) and the JZ, the two functional zones of the mouse placenta, decreased in the L-Tol group (Fig. 1G). A series of mouse blood pressure measurements were recorded during gestation and displayed no significant difference between the L-Tol and the control group (Fig. 1H). These data indicated that the administration of high-dose tolcapone in early gestation caused abortion in mice, whereas the low-dose caused fetal growth restriction in mice with placental damage.

**Figure 1.**
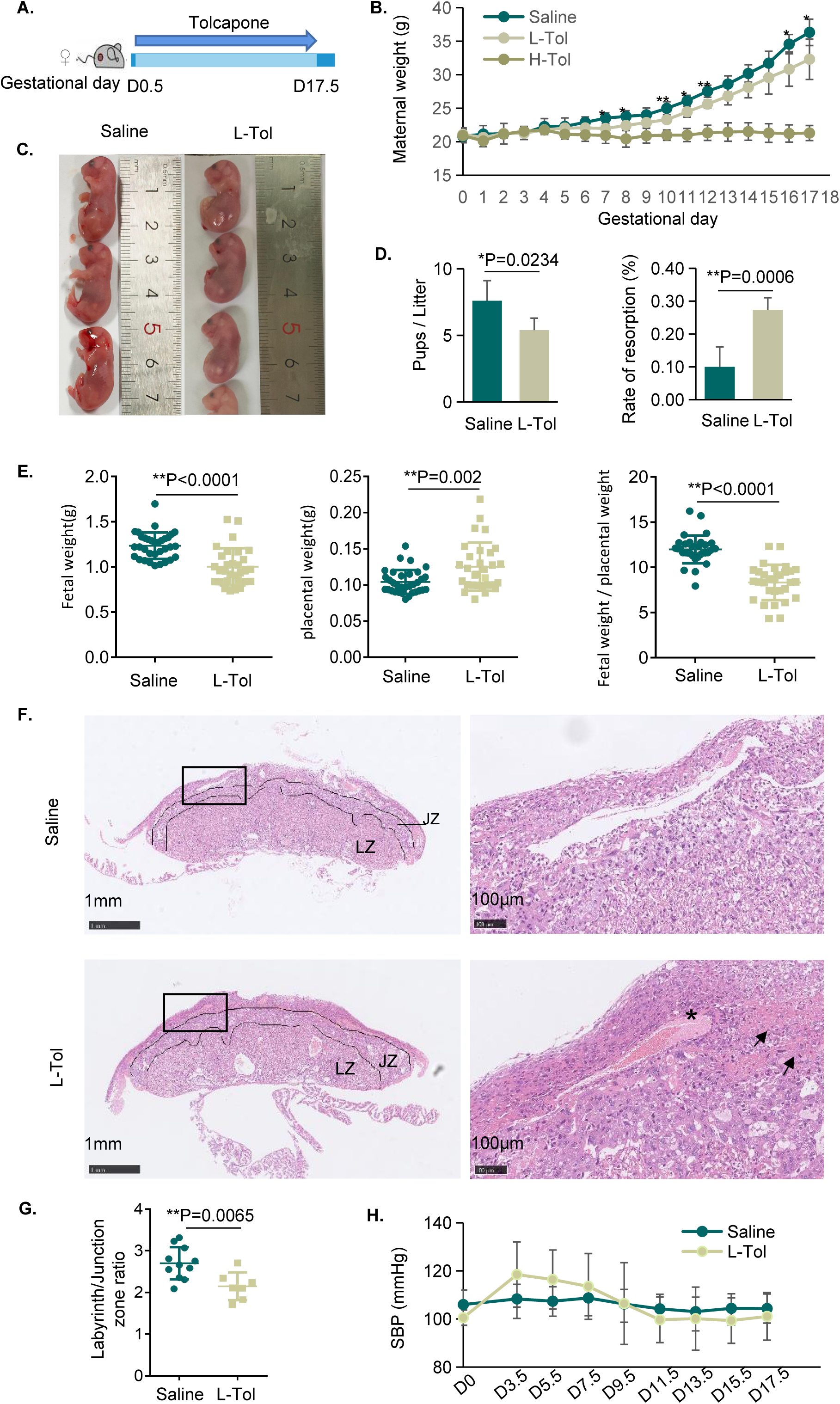
Administration of tolcapone at early gestation resulted in abortion, fetal growth restriction, and placental damage. **A**. Schematic representation of administration of tolcapone. Tolcapone was applied by i.h. injection for 17 days starting at gestational Day 0.5. **B.** Maternal weight during pregnancy with or without tolcapone. H-Tol, high-dose tolcapone; L-Tol, low-dose tolcapone; * indicates p<0.05 in a comparison of saline control group with L-Tol group; ** indicates p<0.01 in a comparison of saline control group with L-Tol group. **C.** Smaller fetal size was observed in the mice with L-Tol comparing with that in the saline control group. **D.** Number of live pups and fetal resorption rate per litter. **E.** Fetal weight and placental weight in mice with or without tolcapone. **F.** Representative H&E staining images of placental tissue sections. Labyrinth zone (LZ) and junction zone (JZ) were indicated by dotted line. The right panel images were zoomed from the left panel showing increased fibrin deposit (marked with arrows) and vascular thrombus formation (marked with a star) in the placentas of L-Tol group. **G.** Ratio of the areas of labyrinth zone over junction zone in placentas. **H.** Maternal systolic blood pressure (SBP) of mice with or without tolcapone treatment was measured during the whole gestation.

### Tolcapone causes preeclampsia-like phenotypes in mid-gestation

Considering the developmental toxicity of tolcapone during early gestation, we examined the effects of tolcapone administration from the middle to the end of gestation. Pregnant mice received high– or low-dose tolcapone starting on gestational day 9.5 and lasting until day 17.5 (Fig. 2A). Maternal body weight increased consistently and showed no significant differences among the three groups (Fig. 2B). Meanwhile, the number of live pups and the fetal resorption rate showed no significant difference with or without tolcapone treatment (Fig. 2E, F), indicating that administration of tolcapone at mid-gestation did not affect mouse abortion, even at a high dose of 60 mg/kg/day. However, tolcapone treatment caused fetal growth restriction even during mid-gestation, as evidenced by decreased fetal size and weight (Fig. 2C, G). Fetal malformations were observed in H-Tol mice (Fig. 2D). Placental weight did not differ with tolcapone treatment (Fig. 2G); however, the morphology of the placentas changed significantly in response to H-Tol and L-Tol. Spongiotrophoblasts, the main trophoblastic cell type comprising of a junction zone, spread irregularly into the labyrinth zone in the placentas of tolcapone-treated mice (Fig. 2H). In addition, pathological changes in the placenta caused the ratios of the two functional zones to decrease (Fig. 2I).

**Figure 2.**
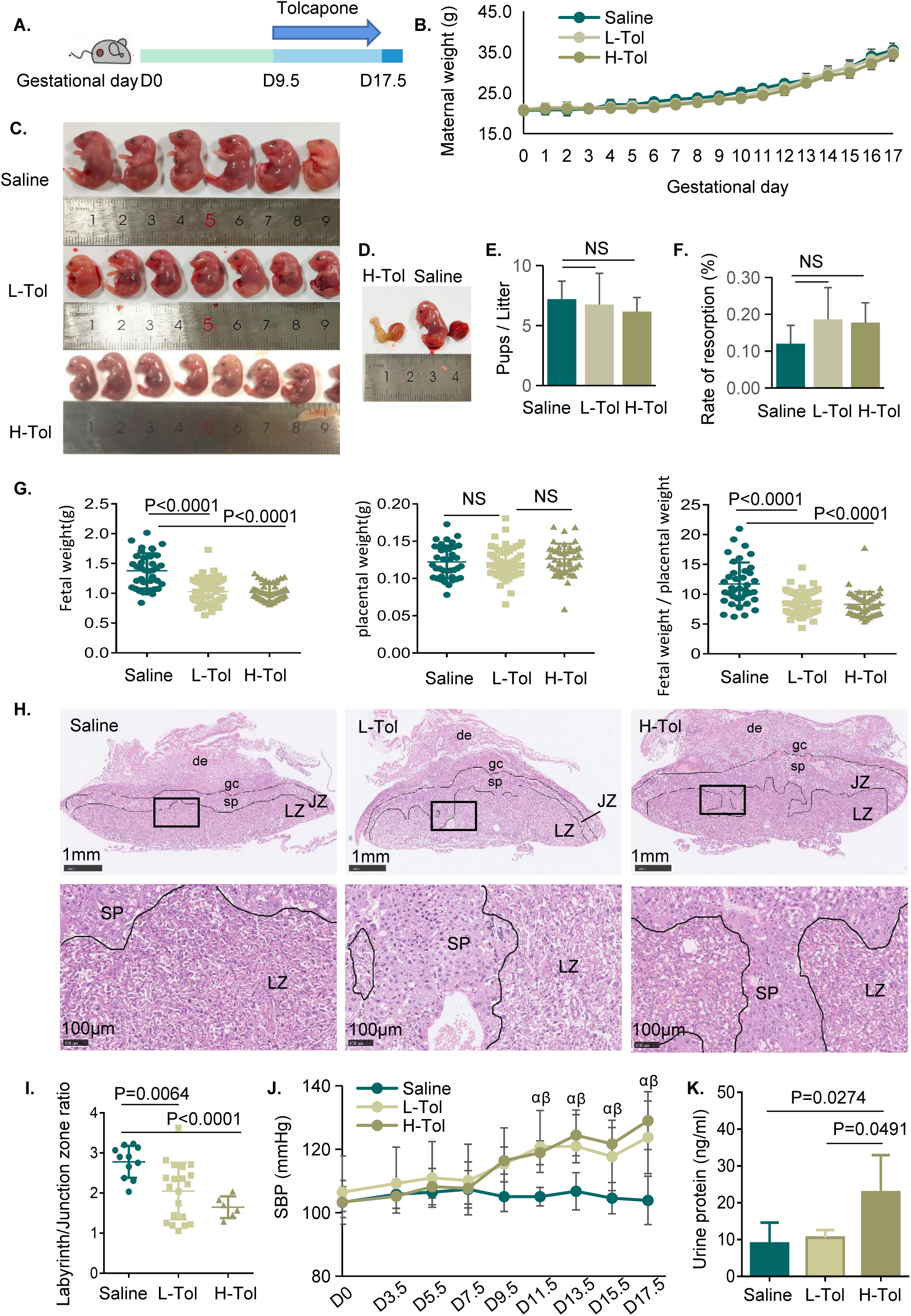
Administration of tolcapone at mid-gestation resulted in fetal growth restriction, maternal gestational hypertension, proteinuria, and placental abnormalities. **A**. Schematic representation of administration of tolcapone. Tolcapone was applied by i.h. injection starting at gestational Day 9.5 until Day 17.5. **B.** Maternal weight during pregnancy with or without tolcapone. **C.** Both L-Tol mice and H-Tol mice showed smaller fetal size when compared with that in the saline control group. **D.** Malformation of embryo from H-Tol mice. **E.** Number of live pups per litter. **F.** Quantification of fetal resorption rate per litter. **G.** Fetal weight and placental weight in the mice of the three groups. **H.** Representative H&E staining images of placental tissue sections. Labyrinth zone (LZ) and junction zone (JZ) were indicated by dotted line. The penetration of spongiotrophoblast cells (SP) in LZ were shown in the placentas of L-Tol and H-Tol mice at the lower panel. de, decidua; gc, trophoblast giant cell. **I.** Ratio of areas of labyrinth zone over junction zone of placentas. **J.** Maternal systolic blood pressure (SBP) of mice was evaluated among the three groups during the whole gestation. α indicates p<0.05 in a comparison of saline group with L-Tol group; β indicates p<0.05 in a comparison of saline group with H-Tol group. **K.** Urinary protein concentration of maternal mice was examined among the three groups at gestational day 17.5.

Maternal blood pressure was examined to determine the function of tolcapone in mothers. Notably, both the H-Tol and L-Tol groups exhibited significantly higher systolic blood pressure (SBP) from gestational day 11.5 to the end of gestation (Fig. 2J). Moreover, H-Tol mice exhibited a significantly higher urine protein concentration on gestational day 17.5, whereas L-Tol did not affect urinary protein excretion (Fig. 2K). These results indicated that tolcapone in mid-gestation led to fetal growth restriction and maternal gestational hypertension, and might be associated with proteinuria if high-dose tolcapone was administered. These symptoms are similar to the typical phenotypes of preeclampsia.

### Tolcapone inhibits the invasion of placental trophoblast to maternal decidua in mice

The invasion of trophoblasts from the placenta to the maternal decidua is critical for placentation and spiral artery transformation[12]. Therefore, we investigated trophoblast invasion in the placental and decidual tissues of mice treated with tolcapone. Immunostainings of cytokeratin 8 (CK8) and α-smooth muscle actin (α-SMA) were conducted in consecutive sections of placental tissue to label placental trophoblasts and maternal decidua, respectively (Fig. 3A). Compared with the saline control, L-Tol mouse placentas showed a significant decrease in trophoblast invasion. The invasion area was further reduced in H-Tol mouse placentas (Fig. 3A, B). In addition to the histomorphological evidence, the EMT marker E-cadherin protein expression was increased in H-Tol mouse placentas (Fig. 3C). These results suggested that tolcapone inhibited trophoblast invasion *in vivo*, potentially through E-cadherin-mediated EMT. To test this hypothesis, we explored the effects of tolcapone on the trophoblast HTR8/SVneo cell line.

**Figure 3.**
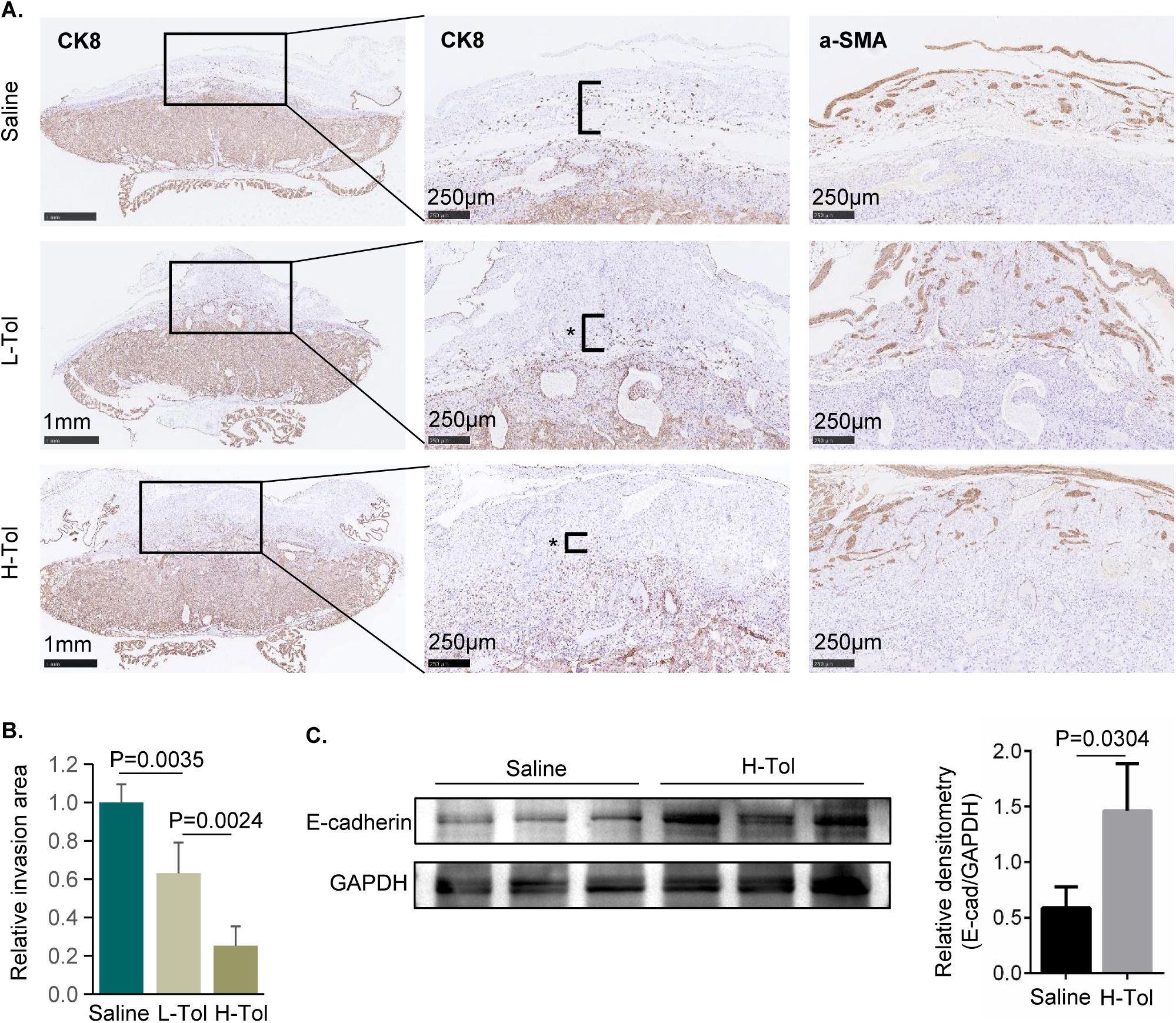
Tolcapone repressed placental trophoblast invasion in mice. **A**. Representative images for immunostainings of cytokeratin 8 (CK8) and α-smooth muscle actin (α-SMA) in consecutive sections of placental tissues. Stars indicated the invasion distance of trophoblasts (CK8 positive cells) in maternal decidua (α-SMA labeled zone). **B.** Relative area of trophoblast invasion into decidua was quantified. **C.** Protein expression level of E-cadherin was detected in placental tissues.

### Tolcapone represses cell invasion in HTR8/SVneo trophoblast cell line

HTR8/SVneo trophoblasts were exposed to different tolcapone concentrations, and cell viability was examined (Fig. 4A). Based on the cell viability curve, we investigated the cell invasion and migration at tolcapone concentrations of 25mM and 35mM. In our study, tolcapone suppressed the migration of HTR8/SVneo cells in a dose– and time-dependent manner (Fig. 4B-D). We further investigated the invasive ability of HTR8/SVneo cells. Tolcapone inhibited cell invasion in the Matrigel-coated Transwells (Fig. 4E). The EMT signaling pathway is a classic regulator of trophoblast invasion and migration. Thus, we explored the potential underlying mechanism by examining Snail, Twist, and E-cadherin protein expressions. The tolcapone-induced inhibition of cell invasion was associated with increased E-cadherin expression and decreased Snail and Twist expressions (Fig. 4F). These data were consistent with observations in mouse placentas and revealed that EMT regulation might be involved in tolcapone-repressed trophoblastic invasion and migration.

**Figure 4.**
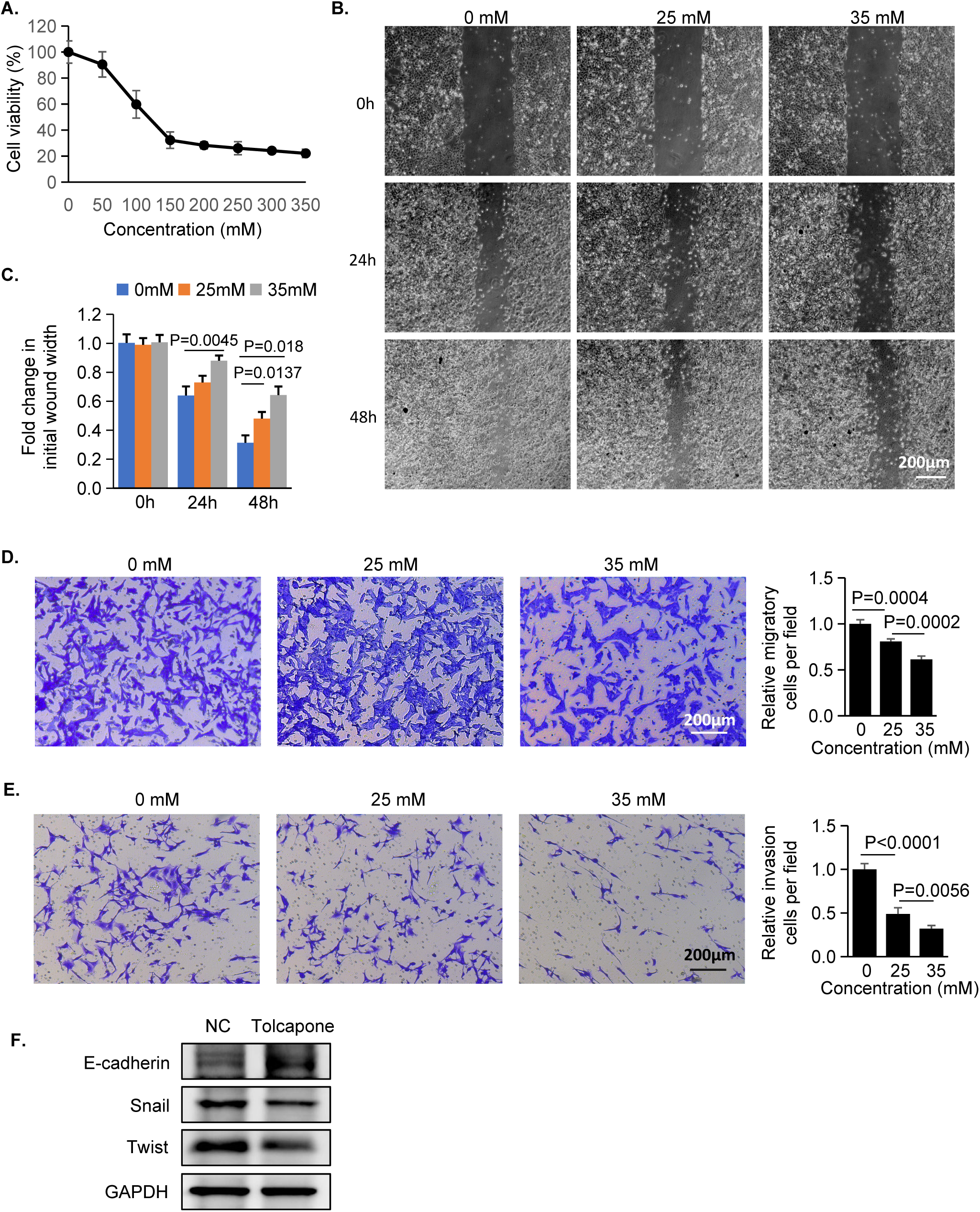
Tolcapone inhibited cell invasion in HTR8/SVneo cells. **A**. Cell viability assay. HTR8/SVneo cells were treated with various doses of tolcapone for 48 h. Cell viability was measured by CCK8 assay. **B.** Wound healing assay of HTR8/SVneo cells treated with 25mM or 35mM tolcapone for 48 h. **C.** Quantification of relative healing wound width is shown. **D.** Measurement of cell migration by Transwell assay. HTR8/SVneo cells were treated with 25mM or 35mM tolcapone for 24 h. **E.** Measurement of cell invasion by Transwell assay. HTR8/SVneo cells treated with 25mM or 35mM tolcapone for 24 h. **F.** Protein expressions of E-cadherin, Snail, and Twist were detected in cells treated with 35mM tolcapone.

### Low concentration of tolcapone induces proliferation in HTR8/SVneo and BeWo trophoblast cell lines

We next determined whether the tolcapone-induced inhibition of cell invasion was followed by the suppression of cell proliferation. HTR8/SVneo cells were treated with 0 mM to 45 mM tolcapone for 72 h and analyzed using cell viability assays. Notably, low concentrations of 5 mM, 15 mM, and 25 mM tolcapone showed proliferation activities, whereas 35 mM tolcapone had no effect on cell viability and 45 mM tolcapone showed evident growth-suppressive activity (Fig. 5A). HTR8/SVneo cells displayed active proliferation when exposed to 25 mM tolcapone (Fig. 5A). To confirm these findings, we performed BrdU staining of HTR8/SVneo cells and observed increased BrdU incorporation into the proliferating nucleus after treatment with 25 mM tolcapone (Fig. 5B). We also tested the effects of tolcapone in a different trophoblast cell line, BeWo. Low concentrations of tolcapone also induced the proliferation of BeWo cells (Fig. 5C).

**Figure 5.**
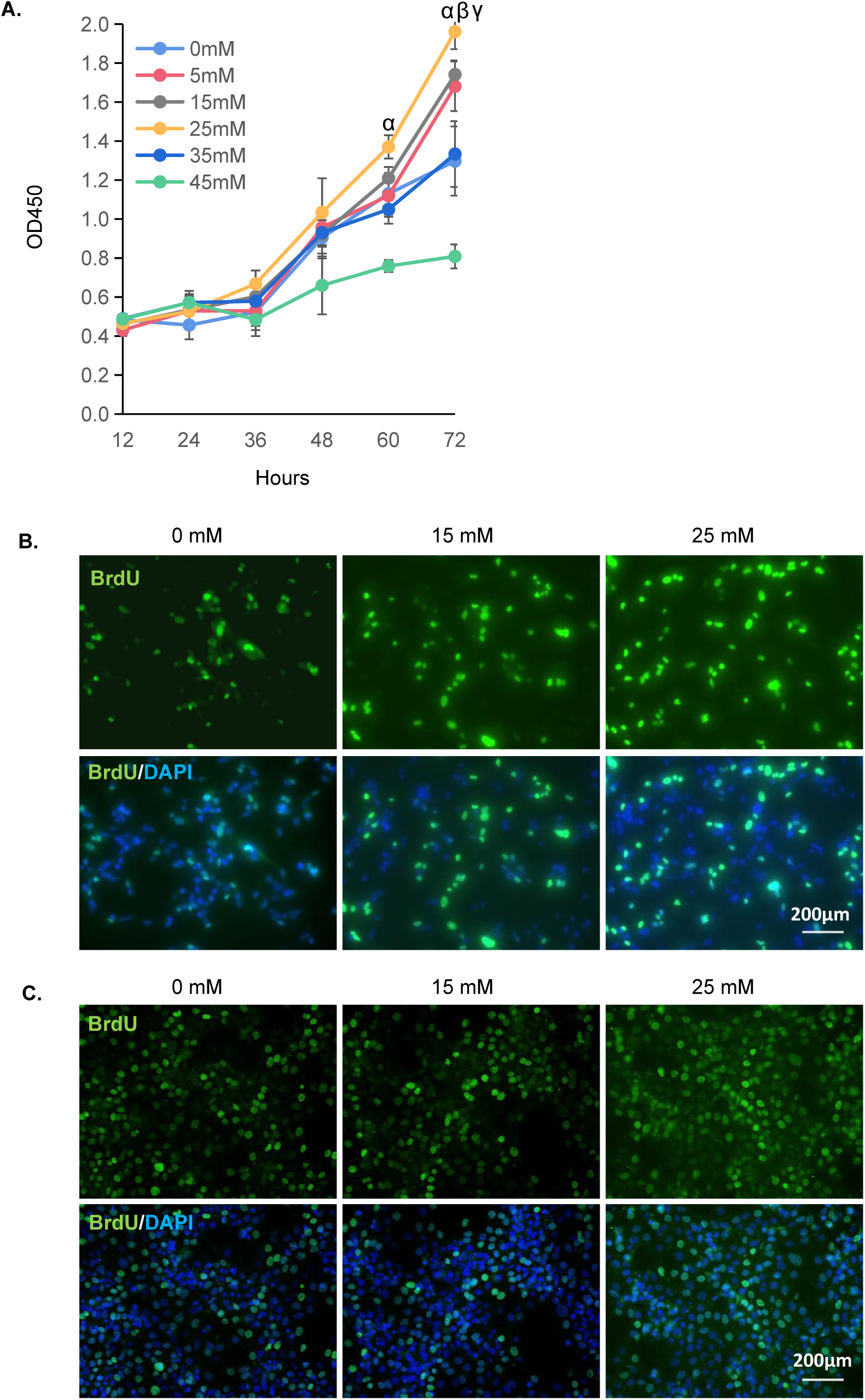
Tolcapone induced proliferation in trophoblasts. **A**. Cell viability assay for the HTR8/SVneo cells treated with various doses of tolcapone for 72 h. α indicates p<0.05 in a comparison of 25 mM group with 0 mM group; β indicates p<0.05 in a comparison of 5 mM group with 0 mM group; γ indicates p<0.05 in a comparison of 15 mM group with 0 mM group. **B.** Representative images of BrdU stained HTR8/SVneo cells treated with tolcapone. **C.** Representative images of BrdU stained BeWo cells treated with tolcapone.

## Discussion

Pregnancy in YOPD is rare, and the clinical experience of caring for pregnant women with PD is limited[20]. However, as the reproductive age increases, it may become a common occurrence[4]. Tolcapone is an effective antiparkinsonian drug used in combination with levodopa/carbidopa. Safety studies of tolcapone in humans and animals have focused on its potential hepatotoxicity; however, the risks to the mother and fetus have not been fully elucidated[10, 21]. Our study demonstrated that tolcapone restricts fetal development and embryo death during early gestation in a dose-dependent manner. The embryo survived if the mother took tolcapone during mid-gestation; however, preeclampsia-like phenotypes associated with fetal growth restriction occurred. One possible mechanism involves EMT signaling-modulated trophoblast invasive dysfunction. Our study provides additional information on the use of tolcapone in pregnancy, including dosage, gestational stage, and adverse effects on the mother and fetus.

Patients with YOPD are more predisposed to levodopa-induced dyskinesias (motor fluctuations)[22]. To overcome the loss of levodopa, two COMT inhibitors, entacapone and tolcapone, are widely used as adjunctive therapies for PD[6]. Based on therapeutic evidence in PD patients with motor fluctuations, tolcapone displays greater efficacy than entacapone[6]. However, evidence regarding the use of tolcapone during pregnancy in women with PD is limited; nonetheless, entacapone has been used in some patients with PD during pregnancy[9, 20, 23]. Animal studies have suggested that entacapone has teratogenic effects on fetal development[24]. Based on these results, women with PD were administered entacapone combined with levodopa/carbidopa starting at week 21 of gestation after the fetal organogenesis period and consequently delivered healthy babies[9, 23]. Our study suggested that tolcapone caused fetal malformation and delayed development in mice, depending on the dosage and gestational stage. Our data showed that low-dose tolcapone in mid-gestation might be safer for fetal formation, although smaller fetuses were observed. In addition, tolcapone induces hypertension in pregnancy, which has not been reported in studies on antiparkinsonian drugs. Maternal preeclampsia-like complications should receive special attention in future studies.

Preeclampsia (PE) is a pregnancy-associated disorder that occures after 20 weeks of gestation with new-onset hypertension, proteinuria, and fetal intrauterine growth restriction[25]. It is widely accepted that preeclampsia is caused by shallow trophoblast invasion. Failure of trophoblast invasion into decidual arteries causes placental hypoxia and releases anti-angiogenic cytokines into the maternal blood, and consequently resulting in maternal hypertension[16, 26]. We observed preeclampsia-like phenotypes in tolcapone-treated mice. Tolcapone administration began at mid-gestation, which is the period of trophoblast invasion, and hampered trophoblast invasion into the placenta and decidua. The *in vitro* experiments verified that tolcapone inhibits the migration and invasion of trophoblasts through changes in EMT signaling. Consistently, excessive E-cadherin hampered trophoblast invasion and has been implicated in patients with preeclampsia[15, 17]. These findings suggest that tolcapone-induced productive and developmental toxicity share a similar mechanism of preeclampsia etiology.

Changes in placental morphology are common in pregnant disorders. Severe placental defects were found in 68% of the embryonic lethal genetic mutant mice[27]. The mutant gene in trophoblast cells caused a higher incidence of embryonic lethality[27]. We noted the abnormal spreading of spongiotrophoblasts in the placentas of tolcapone-treated mice. Consistent with our observations, the infiltration of spongiotrophoblast cells in the labyrinth zone was also detected in *Chtop*−/− and *Pth1r*-/− placentas with obvious fetal developmental delays[27, 28], and to date, its pathogenesis is unknown. The biofunction of the spongiotrophoblast layer is poorly understood; however, it could be a supportive structure of the labyrinth that changes the vessels in the placenta and modulates exchange between the mother and fetus[29].

In our study, the data indicated that tolcapone promotes the proliferation of HTR8/SVneo and BeWo cells. One possible explanation is that the hyperactivated proliferation of trophoblasts results in an increased number of spongiotrophoblasts. However, detecting proliferating trophoblasts in mouse placentas is necessary to validate this hypothesis.

## Conclusion

In the present study, we confirmed that tolcapone caused abortion and fetal growth restriction, and uncovered that tolcapone resulted in maternal hypertension and proteinuria in mice, depending on the dosage and gestational stage at which they received tolcapone. Tolcapone inhibited the invasion of placental trophoblasts *in vivo* and *in vitro*, which was attributed to the regulation of the Snail/Twist/E-cadherin signaling pathway. Our study provides information on using tolcapone in pregnancy with PD, and provides novel insights into the pathogenesis of tolcapone-mediated developmental toxicity.

## Author Contributions

L.C., M.P., Q. J. and F. S. performed experiments and analyzed data. X.Y., Y. M., L. H. designed research and wrote the manuscript. All the authors have read and approved the final version of the manuscript.

## Abbreviations

YOPD: young onset Parkinson’s disease
PD: Parkinson’s disease
COMT: catechol O-methyltransferase
Tol: tolcapone
H-Tol: high-dose tolcapone
L-Tol: low-dose tolcapone
JZ: junction zone
LZ: labyrinth zone
SP: spongiotrophoblast
SBP: systolic blood pressure
DE: decidua
GC: trophoblast giant cell
CK8: cytokeratin 8
α-SMA: α-smooth muscle actin
EMT: epithelial–mesenchymal transition

